# Within-host diversity improves phylogenetic and transmission reconstruction of SARS-CoV-2 outbreaks

**DOI:** 10.1101/2022.06.07.495142

**Authors:** Arturo Torres Ortiz, Michelle Kendall, Nathaniel Storey, James Hatcher, Helen Dunn, Sunando Roy, Rachel Williams, Charlotte Williams, Richard A. Goldstein, Xavier Didelot, Kathryn Harris, Judith Breuer, Louis Grandjean

## Abstract

Accurate inference of who infected whom in an infectious disease outbreak is critical for the delivery of effective infection prevention and control. The increased resolution of pathogen whole-genome sequencing has significantly improved our ability to infer transmission events. Despite this, transmission inference often remains limited by the lack of genomic variation between the source case and infected contacts. Although within-host genetic diversity is common among a wide variety of pathogens, conventional whole-genome sequencing phylogenetic approaches to reconstruct outbreaks exclusively use consensus sequences, which consider only the most prevalent nucleotide at each position and therefore fail to capture low frequency variation within samples. We hypothesized that including within-sample variation in a phylogenetic model would help to identify who infected whom in instances in which this was previously impossible. Using whole-genome sequences from SARS-CoV-2 multi-institutional outbreaks as an example, we show how within-sample diversity is stable among repeated serial samples from the same host, is transmitted between those cases with known epidemiological links, and how this improves phylogenetic inference and our understanding of who infected whom. Our technique is applicable to other infectious diseases and has immediate clinical utility in infection prevention and control.

## Introduction

Understanding who infects whom in an infectious disease outbreak is a key component of infection prevention and control [1]. The use of whole-genome sequencing allows for detailed investigation of disease outbreaks, but the limited genetic diversity of many pathogens often hinders our understanding of transmission events [2]. As a consequence of the limited diversity, many index case and contact pairs will share identical genotypes, making it difficult to ascertain who infected whom.

Within-sample genetic diversity is common among a wide variety of pathogens [3–7]. This diversity may be generated *de novo* during infection, by a single transmission event of a diverse inoculum or by independent transmission events from multiple sources [8]. The maintenance and dynamic of within-host diversity is then a product of natural selection, genetic drift, and fluctuating population size [1]. The transmission of within-host variation between individuals is also favored as a large inoculum exposure is more likely to give rise to infection [9–14].

Most genomic and phylogenetic workflows involve either genome assembly or alignment of sequencing reads to a reference genome. In both cases, conventionally the resulting alignment exclusively represents the most common nucleotide at each position. This is often referred to as the consensus sequence. Although genome assemblers may output contigs (combined overlapping reads) representing low frequency haplotypes, only the majority contig is kept in the final sequence. In a mapping approach, a frequency threshold for the major variant is usually pre-determined, under which a position is considered ambiguous. The lack of genetic variation between temporally proximate samples and the slow mutation rate of many pathogens results in direct transmission events sharing exact sequences between the hosts when using the consensus sequence approach. For instance, the substitution rate of SARS-CoV-2 has been inferred to be around 2 mutations per genome per month [15]. Given its infectious period of 6 days [16], most consensus sequences in a small-scale outbreak will show no variation between them. This lack of resolution and poor phylogenetic signal complicate the determination of who infected whom, relying exclusively on epidemiological information.

We hypothesize that the failure of consensus sequence approaches to capture within-sample variation arbitrarily excludes meaningful data and limits the ability to determine who infected whom, and that including within-sample diversity in a phylogenetic model would significantly increase the evolutionary and temporal signal and thereby improve our ability to infer infectious diseases transmission events. We tested our hypothesis on multi-institutional SARS-CoV-2 outbreaks across London hospitals that were part of the COVID-19 Genomics UK (COG-UK) consortia [17]. Technical replicates, repeated longitudinal sampling from the same patient, and epidemiological data allowed us to evaluate the presence and stability of within-sample diversity within the host and in independently determined transmission chains. We also evaluated the use of within-sample diversity in phylogenetic analysis by conducting simulations of sequencing data using a phylogenetic model that accounts for the presence and transmission of within- sample variation. We show the effects on phylogenetic inference of using consensus sequences in the presence of within-sample diversity, and propose that existing phylogenetic models can leverage the additional diversity given by the within-sample variation and reconstruct the phylogenetic relationship between isolates. Lastly, we show that by taking into account within-sample diversity in a phylogenetic model we improve the temporal signal in SARS- CoV-2 outbreak analysis. Using both phylogenetic outbreak reconstruction and simulation we show that our approach is superior to the current gold standard whole-genome consensus sequence methods.

## Results

### Sampling, demographics and metadata

Between March 2020 and November 2020, 451 healthcare workers, patients and patient contacts at the participating North London Hospitals were diagnosed at the Camelia Botnar Laboratories with SARS-CoV-2 by PCR as part of a routine staff diagnostic service at Great Ormond Street Hospital NHS Foundation Trust (GOSH). The mean participant age was 40 years old (median 38.5 years old, interquartile range (IQR) 30-50 years old), and 60% of the participants were female (Supplementary Table 1). A total of 289 were whole-genome sequenced using the Illumina NextSeq platform, which resulted in 522 whole-genome sequences including longitudinal and technical replicates (Supplementary Data File 1). All samples were SARS-CoV-2 positive with real time qPCR cycle threshold (C*_t_*) values ranging from 16 to 35 cycles (Supplementary Table 1). The earliest sample was collected on 26th March 2020, while the latest one dated to 4th November 2020 (Supplementary Fig. 1a). A total of 291 samples had self-reported symptom onset data, for which the mean time from symptom onset to sample collection date was 5 days (IQR 2-7 days, Supplementary Fig. 1b). More than 90% of the samples were taken from hospital staff, while the rest comprised patients and contacts of either the patients or the staff members (Supplementary Table 1).

### Genomic analysis of SARS-CoV-2 sequences

Whole-genome sequences were mapped to the reference genome resulting in a mean coverage depth of 2177x (Supplementary Fig. 2). A total of 454 whole-genomes with mean coverage higher than 10x were kept for further analysis. Allele frequencies were extracted using the pileup functionality within *bcftools* [18] with a minimum base and mapping quality of 30, which represents a base call error rate of 0.1%. Variants were filtered further for read position bias and strand bias. Only minor variants with an allele frequency of at least 1% were kept as putative variants. Samples with a frequency of missing bases higher than 10% were excluded, keeping 350 isolates for analysis. The mean number of low frequency variants was 12 (median 3, IQR 1.00 – 9.75), although both the number of variants and its deviation increased at high C*_t_* values (Supplementary Fig. 3).

### Within-sample variation is stable between technical replicates

To understand the stability of within-sample variation and minimize spurious variant calls, we sequenced and analyzed technical replicates of 17 samples. Overall, when the variant was present in both duplicates the correlation of the variant frequencies was high (R^2^ = 0.9, Fig. 1a right). The high correlation was also maintained at low variant frequencies (Fig. 1a left).

**Figure 1:**
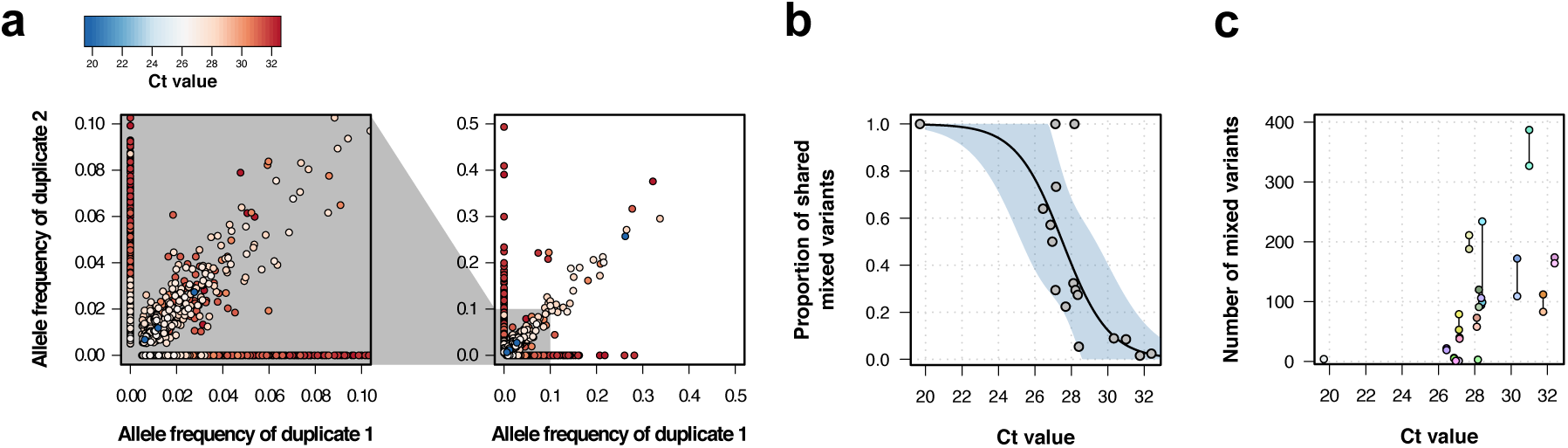
Genomic analysis of technical duplicates. a. Allele frequency comparison between technical replicates for all frequencies (right) and for frequencies up to 1% (left). Colors represent the C*_t_* value for the sample. **b** Proportion of shared mixed variants between technical replicates in relation to the C*_t_* value. **c** Total number of mixed variants in relation to the C*_t_* value. Lines linked two technical replicates. Each sequence has a different color, with sequences from the same patient having a different shade of the same color.

Minor variants were less likely to be detected or shared when one or more of the paired samples had a low viral load. These discrepancies may appear because of amplification bias caused by low genetic material, base calling errors due to low coverage, or low base quality. The mean proportion of discrepant within-sample variants between duplicated samples was 0.39 (sd = 0.29), although this varied between duplicates (Supplementary Fig. 4). C*_t_* values in RT-PCR obtained during viral amplification are inversely correlated with low viral load [19]. The proportion of shared intra-host variants was negatively correlated with C*_t_* values in a logistic model (estimate=-0.78, p-value=0.008), with higher C*_t_* values associated with a lower amount of shared intra-host variants (Fig. 1c). The number of within-sample variants detected also increased with C*_t_* value, as well as the deviation in the number of variants between duplicates (Fig. 1d). This could be explained either by an increase in the number of spurious variants at low viral loads [20], biased amplification of low level sub-populations minor rare alleles [21], or due to the accumulation of within-host variation through time, as viral load (with rising C*_t_* values) also decreases from time since infection.

Based on these results, only samples with a C*_t_* value equal or lower than 30 cycles were considered, which resulted in 249 samples kept for analysis. For the filtered dataset, 414 out of 29903 positions were polymorphic for the consensus sequence, while the alignment with within-sample diversity had 1039 SNPs. Of these, 699 positions had intra-host diversity, of which 78% (549/699) were singletons. The majority of samples (207/249, 83%) contained at least 1 position with a high quality within-host variant, and the median amount of intra-host variants per sample was 2 (IQR 1-4.5).

### Within-sample variation is shared between epidemiologically linked samples

Given the limited genomic information in the consensus sequences, epidemiological data is often necessary to infer the directionality of transmission. We performed a pairwise comparison of all samples and calculated the proportion of shared within-sample variants (shared variants divided by total variants in the pair). We compared samples that a) did not have any recorded epidemiological link, b) samples that were from the same hospital (possibly linked), c) samples that were part of the same department within the same hospital (probable link), and d) samples that had an epidemiological link within the same department of the same hospital (proven link), e) were a longitudinal replicate from the same patient and f) a technical replicate from the same sample.

We tested the concordance between epidemiological and genomic data by determining the genetic distance between pairs of samples with epidemiological links and without them. Pairs of samples from the same hospital, department, epidemiologically linked, or longitudinal and technical replicates were more closely located in the consensus phylogenetic tree than those samples that did not have any relationship, although this difference was small in the case of pairs of samples from the same hospital (Table 1).

**Table 1:** Phylogenetic distance (substitutions per genome per year) between pairs of samples.

| Sample relationship | Estimate (95%CI) | p-value |
| --- | --- | --- |
| None | 11.73 (11.63 - 11.83) | Reference |
| Hospital | 10.3 (10.09 - 10.54) | $<1 \times 10^{-4}$ |
| Department | 5.35 (4.21 - 6.49) | $<1 \times 10^{-4}$ |
| Epidemiological | 1.62 (0.53 - 2.72) | $<1 \times 10^{-4}$ |
| Longitudinal duplicates | 0.01 (-1.84 - 1.86) | $<1 \times 10^{-4}$ |
| Technical replicate | 0 (-4.29 - 4.29) | $<1 \times 10^{-4}$ |

The proportion of shared within-host variants was significantly higher between technical replicates, longitudinal duplicates, epidemiologically linked samples, and samples taken from individuals from the same department when compared to pairs with no epidemiological links (Fig. 2). The probability of sharing a low frequency variant was inferred using a logistic regression model (Supplementary Fig. 5). There was a tendency for the probability to increase with variant frequency, but the association was not strong (Odds ratio 1.8, 95% CI 0.9 – 3.5, p=0.08). The probability of sharing a variant for samples with no epidemiological links was 9.5 *×* 10*^−^*^6^ (95% CI 8.8 *×* 10*^−^*^6^ – 1.02 *×* 10*^−^*^5^). Samples from the same hospital did not have a probability significantly higher than those without any link (3.3 *×* 10*^−^*^3^, 95% CI 2.7 *×* 10*^−^*^3^ –4.03 *×* 10*^−^*^3^). On the other hand, pairs from the same department, with epidemiological links, replicates or technical replicates all had a higher probability of sharing a low frequency variant when compared to those pairs with no link (all p-values *<* 0.001). The inferred probabilities for pairs from the sample department was 1.4% (95% CI 0.9% – 2.1%), which increased to 5% for pairs with epidemiological links (95% CI 4.2% – 6.4%). For longitudinal replicates, the probability was inferred to be 38% (95% CI 35% – 41%), while technical replicates were estimated to have the highest probability (70%, 95% CI 64% – 76%).

**Figure 2:**
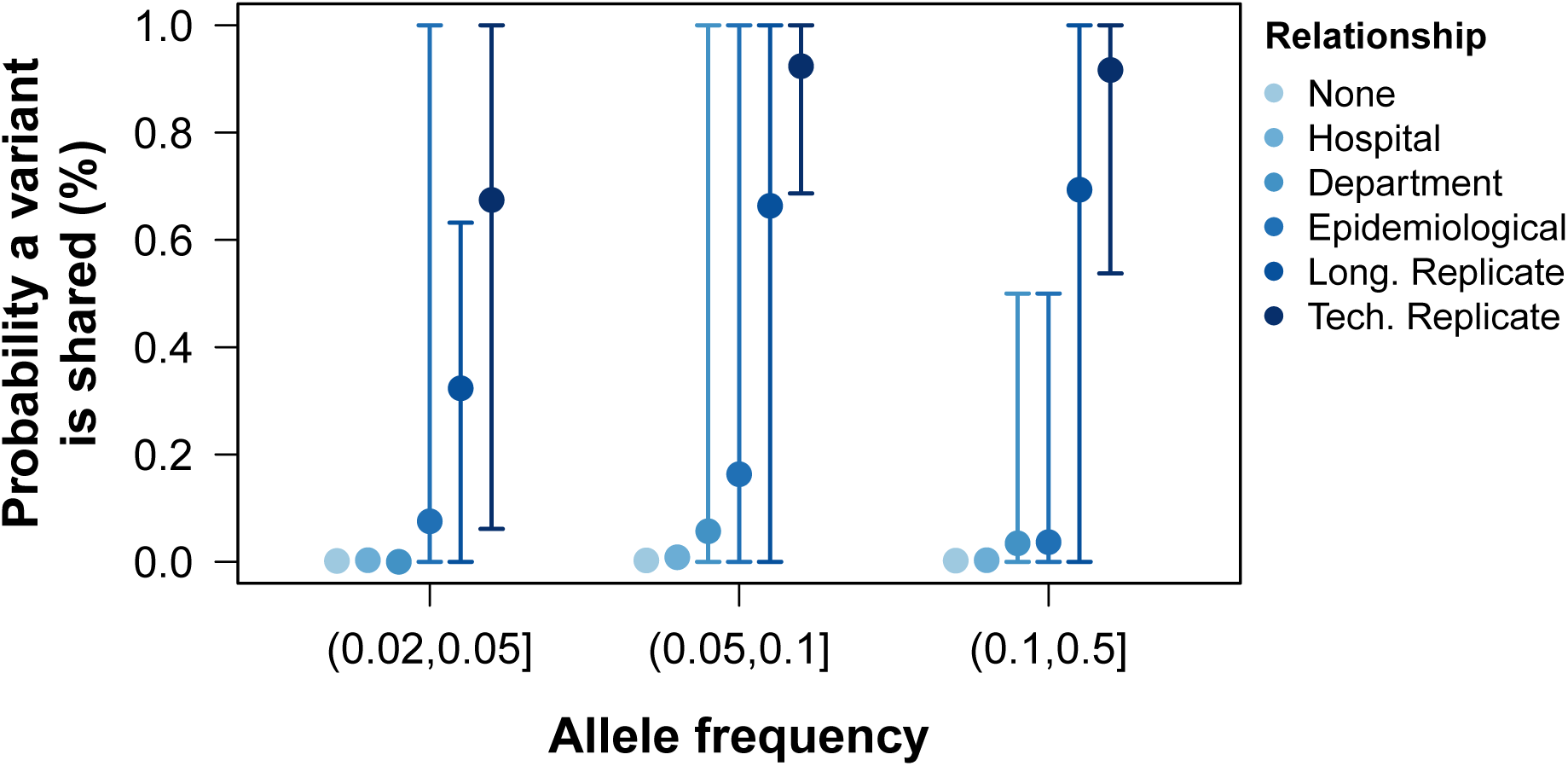
Probability of sharing within-host variants in sample pairs. The probability of variants shared between pairs of samples calculated as the number of low frequency variants in both samples divided by the total number of variants between the pair. Colors grouped samples by their relationship. Points represent the mean probability a variant is shared between all pairwise samples within a group and allele frequency. Error bars show the 95% CI.

### Within-host diversity model outperforms the consensus model in simulations

The effect of within-sample diversity in phylogenetic inference was tested by evaluating the accuracy in the reconstruction of known phylogenetic trees using a conventional phylogenetic model and a model that accounts for within-sample variation.

The presence of within-sample diversity was coded in the genome alignment using existing IUPAC nomenclature [22]. For the consensus sequence alignment, only the 4 canonical nucleotides were used (Fig. 3a,b), while the proposed alignment retained the major and minor allele information as independent character states (Fig. 3c,d).

**Figure 3:**
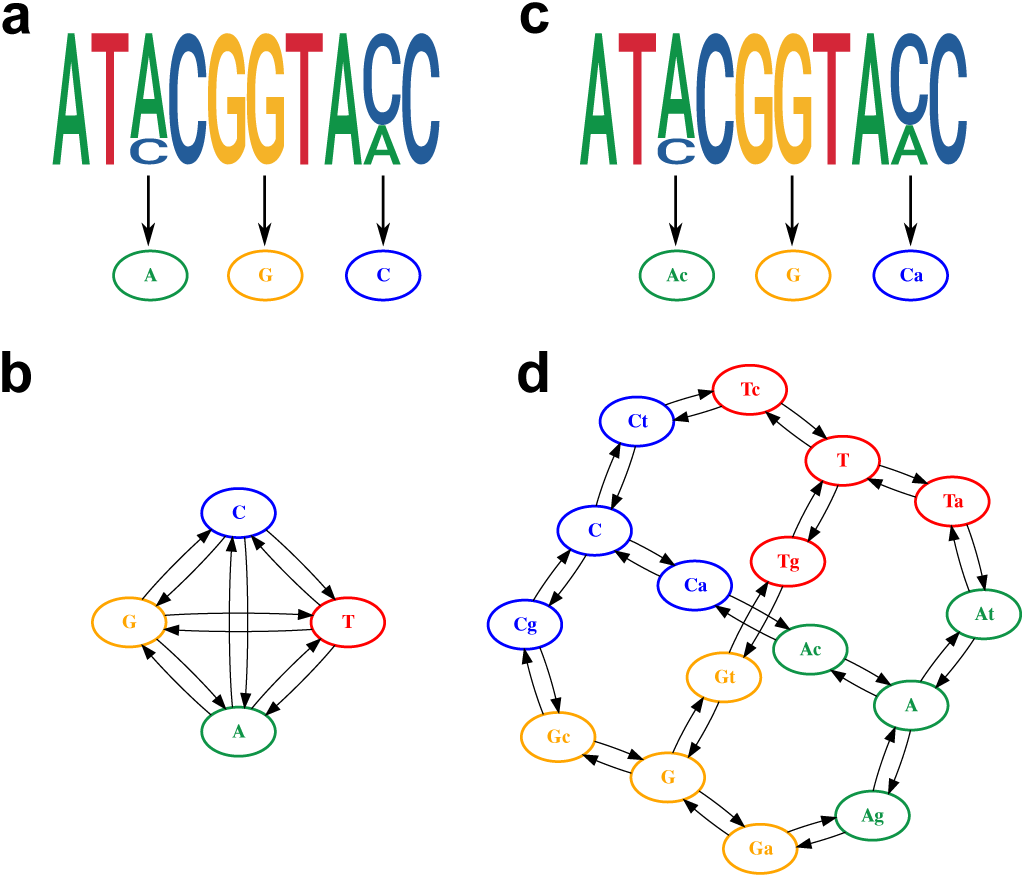
Model of within-host diversity. Proposed evolutionary model of within-host diversity in genomic sequences. Upper- case letters represent the major variant in the population, while lowercase letters indicate presence of a minor variant alongside the major one. **a, c** Genome sequences where some positions show within-sample variation (top), represented by a major allele (big size letter) and a minor one (smaller size), as well as its representation in the alignment (bottom). **b, d** Models of nucleotide evolution. Character transitions are indicated by arrows. **a** Consensus sequence, where only the major allele is represented in the alignment. **b** Model of nucleotide evolution using the consensus sequence, with four character states representing the four nucleotides. **c** Sequence with within-sample variation, represented by an uppercase letter for the major allele and a lower case letter for the minor allele. **d** Model of nucleotide evolution with 16 character states accounting for within-sample variation.

In order to evaluate the bias in tree inference with and without the inclusion of within-sample diversity, we simulated genome alignments for 100 random trees using a phylogenetic model where both major and minor variant combinations were considered, resulting in a total of 16 possible states (Fig. 3d) and the substitution rates shown in Supplementary Table 3. From the simulated genomes, two types of alignments were generated: a consensus sequence, where only the major allele was considered (Fig. 3a); and an alignment that retained the major and minor allele information as independent character states (Fig. 3c). From the simulated alignments, RaxML-NG was used to infer phylogenetic trees [23]. The consensus sequence was analyzed with a GTR+*γ* model, while the PROTGTR+*γ* model was used in order to accommodate the extra characters of the alignment with within-sample diversity and major/minor variant information.

The two models were evaluated for their ability to infer the known phylogeny that included within-host diversity. The estimated phylogenies were compared to the known tree using different measures to capture dissimilarities in a variety of aspects relevant to tree inference (Supplementary Table 2). For all the metrics employed, the phylogenies inferred explicitly using within-host diversity as independent characters approximated better to the initial tree than the one using the consensus sequence (Fig. 4). Additionally, the transition/transversion rates inferred by the phylogenetic models accounting for within-host diversity accurately reflect the rates used for the simulation of genomic sequences (Supplementary Table 3-5).

**Figure 4:**
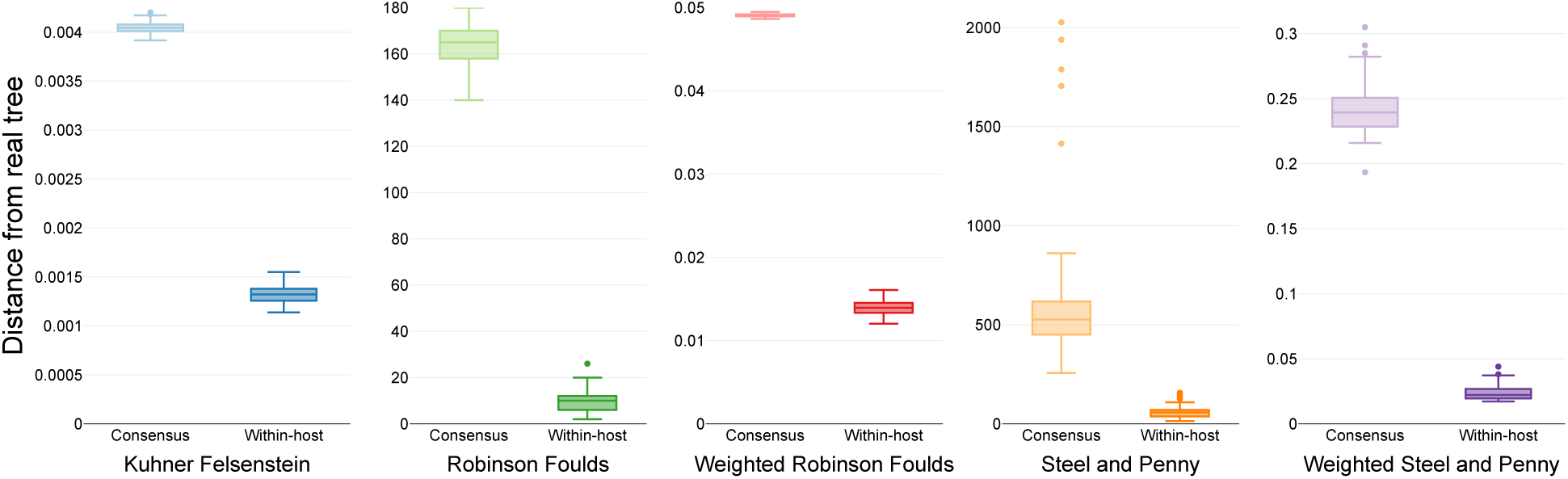
Similarity scores for inferred trees. Comparison of the phylogenetic trees inferred using three simulated sequences and different phylogenetic models with the known starting tree. Colors differentiate the metrics used for the comparison.

### Within-host diversity improves the resolution in SARS-CoV-2 phylogenetics

Genome sequences collected at different time points are expected to diverge as time progresses, resulting in a positive correlation between the isolation date and the number of accumulated mutations (temporal signal) [24]. The alignment with consensus sequences and the one reflecting within-sample variation were used to infer two different phylogenetic trees (Supplementary Fig. 6). Longitudinal samples in the phylogeny inferred using within-host diversity reflected the expected temporal signal, with an increase in genetic distance as time progressed between the longitudinal pairs in a linear model (coefficient 2.24, 0.59 - 3.88 95% CI, p = 0.019, Supplementary Fig. 7). The difference in C*_t_* value among longitudinal duplicates was not correlated with a higher genetic distance (coefficient 1.62, -0.66 - 3.91 95% CI, p = 0.2).

We analyzed the impact of using within-sample variation on the temporal structure of the phylogeny by systematically identifying clusters of tips in the phylogenetic tree with an identical consensus sequence and no temporal signal. We then performed a root-to-tip analysis using the tree inferred with intra-sample diversity. Only clusters with more than 3 tips were used for the root-to-tip analysis. The majority of clusters (10/11) showed a positive correlation between the distance of the tips to the root and the collection dates, demonstrating a significant temporal signal between samples when there was none using the conventional consensus tree (Fig. 5).

**Figure 5:**
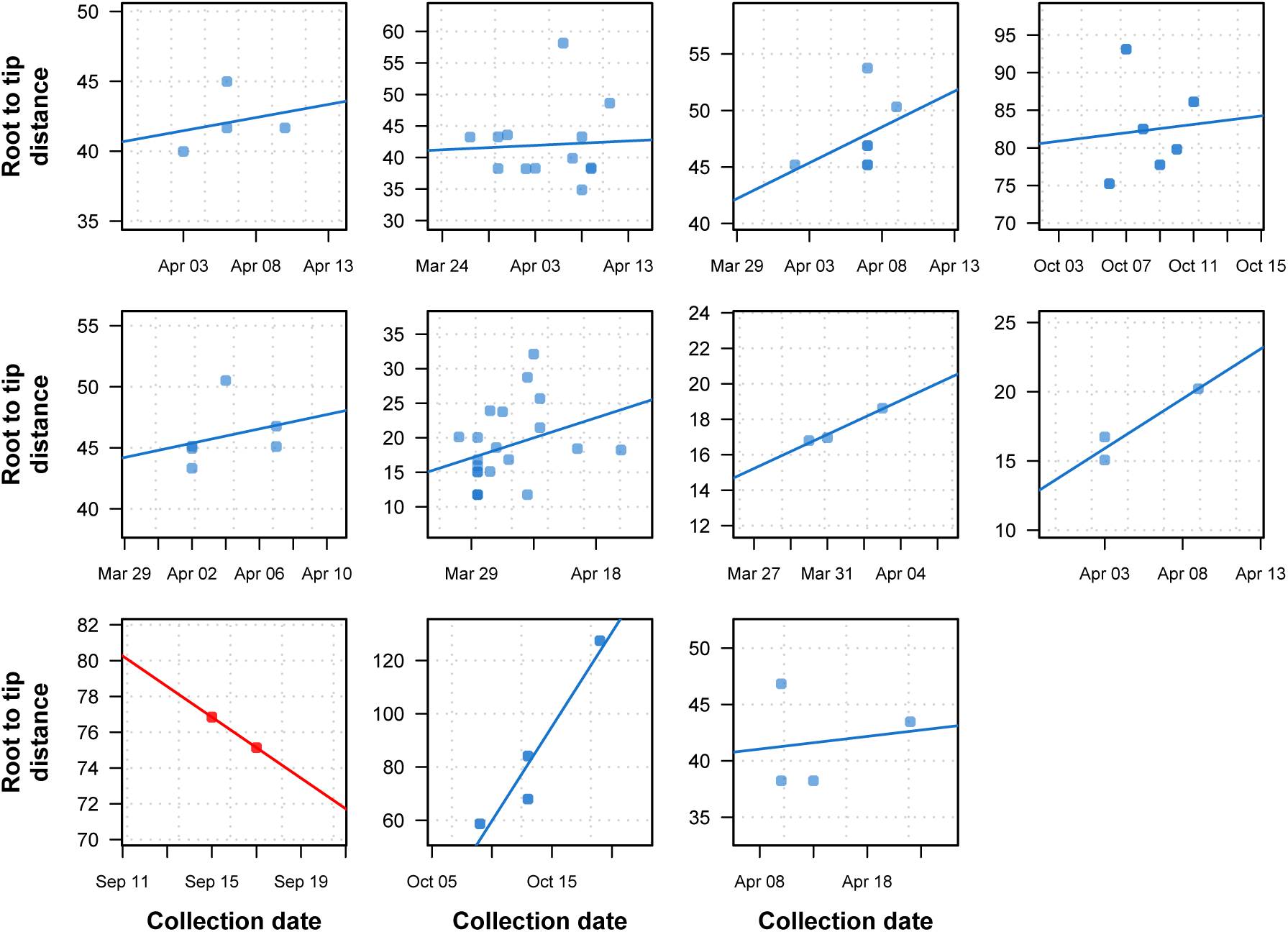
Previously uninformative clusters present temporal signal when using within-sample diversity. A set of 11 outbreak clusters (one per panel, each plotting the root to tip distance against time) in which all samples had identical consensus genomes sequences (and therefore no temporal signal). Blue colors indicate those regressions that after utilizing within sample diversity now have a positive slope (temporal signal), and red shows those regressions that have a negative slope (misleading or false positive temporal signal).

To illustrate the downstream application of the improved phylogenetic resolution, we inferred a time-calibrated phylogeny with the collection dates of the tips using BactDating [25] (Supplementary Fig. 8) and calculated the likelihood of transmission events within potential epidemiologically identified outbreaks using a Susceptible-Exposed-Infectious-Removed (SEIR) model [26]. The SEIR model was parameterized with an average latency period of 5.5 days [27], an infectious period of 6 days [16], and a within-host coalescent rate of 5 days as previously estimated for SARS-CoV-2 [28]. The likelihood of transmission was calculated for every pair of samples, while the Edmonds algorithm as implemented in the R package *RBGL* [29] was used to infer the graph with the optimum branching (Fig. 6c,d; Supplementary Fig. 9).

**Figure 6:**
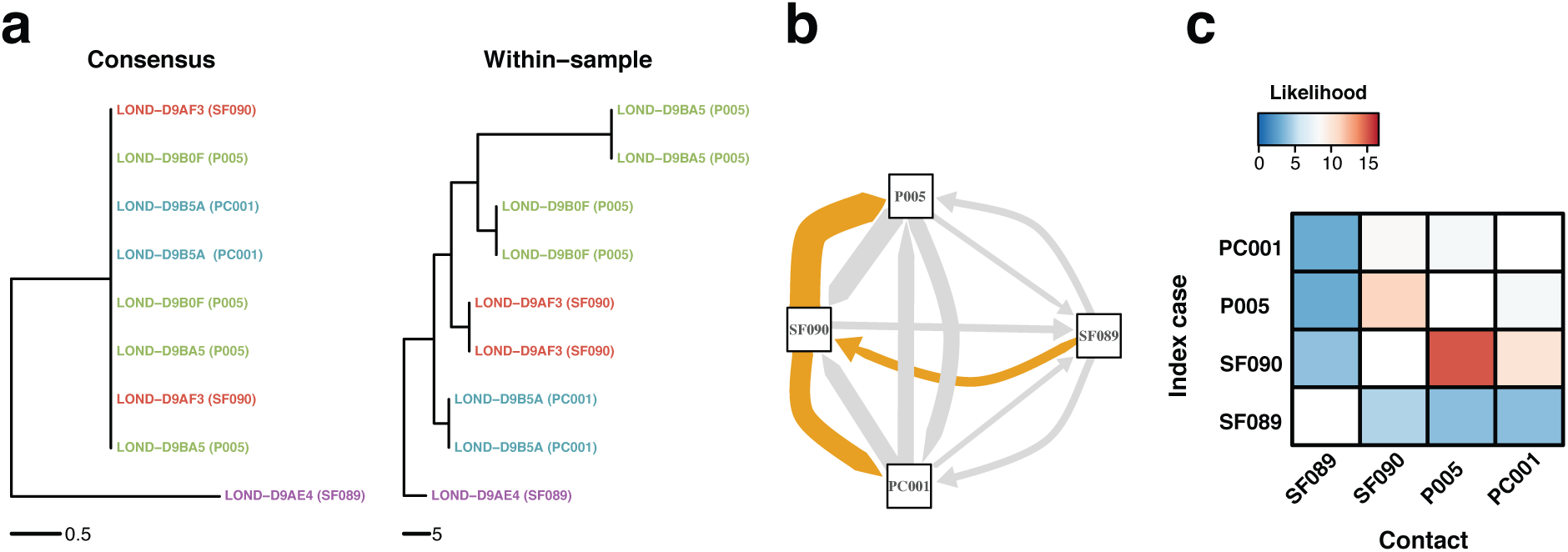
Within-sample variation improves resolution of infectious disease outbreaks. Effect of using low frequency variants in phylogenetic inference. **a** Maximum likelihood phylogeny using the consensus sequences (left) and the alignment leveraging within-sample variation. Replicates of the same sample share the same color. Sample IDs are coded as follows: SF, for staff members; P, for patients; and PC, for patient contacts. **b** Transmission network inferred using within sample variation. Edge width is proportional to the likelihood of direct transmission using a Susceptible-Exposed-Infectious-Removed (SEIR) model. Colored edges represent the Edmunds optimum branching and thus the most likely chain. **c** Heatmap of the likelihood of direct transmission between all pairwise pairs of samples using a SEIR model. Vertical axis is the infector while the horizontal axis shows the infectee.

## Discussion

Detailed investigation of transmission events in an infectious disease outbreak is a prerequisite for effective prevention and control. Although whole-genome sequencing has transformed the field of pathogen genomics, insufficient pathogen genetic diversity between cases in an outbreak limits the ability to infer who infected whom. Using multi-hospital SARS-CoV-2 outbreaks and phylogenetic simulations, we show that including the genetic diversity of sub- populations within a clinical sample improves phylogenetic reconstruction of SARS-CoV-2 outbreaks and determines the direction of transmission when using a consensus sequence approach fails to do so.

The majority of samples sequenced harbored variants at low frequency that remained stable in technical replicates. However, within-host variation was less consistent between paired samples with a lower viral load (higher C*_t_*). This is likely to be a consequence of low starting genetic material giving rise to amplification bias during library preparation and sequencing. Establishing a cut-off for high C*_t_* values is therefore important to accurately characterize within-host variation. In our study, we excluded samples with a C*_t_* value higher than 30 cycles based on the diagnostic PCR used at GOSH. Since C*_t_* values are only a surrogate for viral load and are not standardized across different assays [30], appropriate thresholds would need to be determined for other primary PCR testing assays.

The generation, maintenance and evolution of subpopulations within the host reflect evolutionary processes which are meaningful from phylogenetic and epidemiological perspectives. Subpopulations within a host can emerge from three mechanisms: de novo diversification in the host, transmission of a diverse inoculum, or multiple transmission events from different sources. If the subpopulations are the result of de novo mutations, nucleotide polymorphisms within the subpopulations accumulate over time and may therefore result in a phylogenetic signal useful for phylogenetic inference. In our data, longitudinal samples taken at later time points were demonstrated to accrue genomic variation. Although this pattern can be confounded by decreasing viral load as infection progresses, C*_t_* values in our dataset were not correlated with a higher genetic distance, and clusters in our data containing both longitudinal and technical replicates also corroborate these results. Transmission of a diverse inoculum also gives rise to phylogenetically informative shared low frequency variants, as our results show that immediate transmission pairs are more likely to share variants at low frequency. The effect of multiple transmission events in the phylogeny depends on the relatedness of both index cases and the bottleneck size in each transmission event.

Paired samples with epidemiological links and from the same department shared a higher proportion of low frequency variants and were located closer in the consensus tree than samples with no relationship. Similarly, samples with shorter distance in the consensus phylogeny were more likely to share low frequency variants. These patterns suggest that the distribution of low frequency variants is linked to events of evolutionary and epidemiological interest. The fact that technical duplicates shared more within-host diversity than longitudinal replicates of the same sample suggests that much of the variation within hosts is transitory. Therefore, within-host diversity may be relevant on relatively short time scales, which is precisely where consensus sequences lack resolution. Combining the data derived from fixed alleles in the consensus sequences and transient within-sample minor variation enables an improved understanding of the relatedness of pathogen populations between hosts.

The effects of neglecting within-host diversity in phylogenetic inference were analyzed by using simulated sequences under a phylogenetic model that reflects the presence and evolution of within-host diversity. We compared a conventional consensus phylogenetic model and a model that leverages within-sample diversity, and evaluated their ability to infer the known phylogeny. Our proposed phylogenetic model incorporates within-sample variation by explicitly coding major and minor nucleotides as independent characters in the alignment. We demonstrated that phylogenies inferred using the conventional consensus sequence approach were heavily biased and unrepresentative of the known structure of the simulated tree. However, sequences that included within-host diversity were shown to infer less biased phylogenetic trees.

Previous studies have addressed the use of within-host variation to infer transmission events. Wymant et al. [31] employed a framework based on phylogenetic inference and ancestral state reconstruction of each set of populations detected within read alignments using genomic windows. Our study extends this work by coding genome-wide diversity within the host directly in the alignment and the phylogenetic model. De Maio et al [32] proposed direct inference of transmission from sequencing data alongside host exposure time and sampling date within the bayesian framework BEAST2 [33]. Our approach is focused on directly improving the temporal and phylogenetic signal of whole-genome sequences, and it’s especially suited for use in applications and analysis that employ a phylogenetic tree as input to infer transmission [34].

Future work will extend this model by including allele frequency data in addition to independent characters for major and minor variants. Phylogenetic models that explicitly include dynamics of within-sample variation and sequencing error may further improve phylogenetic inference or allow researchers to better estimate parameters of interest, including R0, bottleneck size, transmissibility and the origin of outbreaks.

Our study benefited from the availability of sequenced technical replicates that enabled us to distinguish genuine variation from sequencing noise, especially at low variant frequencies. Similarly, access to longitudinal samples from the same patient allowed us to characterize the spectrum of within host variation and therefore reconstruct transmission chains with more precision.

In line with conventional consensus sequencing approaches, we used a reference sequence for genome alignment and variant calling. Although widely used, one limitation of this approach is a potential mapping bias causing some reads to reflect the reference base at low frequencies at a position where only a variant should be present. Although we applied stringent quality filtering, we cannot rule out the persistence of some false positive minor variants. Using genome graphs to map to a reference that encompasses a wider spectrum of variation may alleviate this problem, and could be an interesting addition to pathogen population genomic analysis.

Our results demonstrate that within-sample variation can be leveraged to increase the resolution of phylogenetic trees and improve our understanding of who infected whom. Using SARS-CoV-2 as an example, we show that variants at low frequencies are stable, phylogenetically informative and are more often shared among epidemiologically related contacts. We propose that pathogen phylogenetic models should accommodate within-host variation to improve the understanding of infectious disease transmission.

## Materials and methods

### Model for within-host diversity

Whole-genome alignments were generated from 100 random phylogenetic trees with 100 tips with the function *SimSeq* of the R package *phangorn* [35, 36] using a model with 16 character states that represent the combinations of the 4 nucleotides with each other as minor and major alleles (Fig. 3d). Three substitution rates for the model were considered: a rate at which minor variants evolve, equal to 1; the rate at which minor variants are lost, leaving only the major nucleotide at that position, equal to 100; and the rate at which minor/major variants are switched, equal to 200.

Two types of alignments were generated from the simulated genomes: a consensus sequence, where only the major allele was considered; and an alignment that retained the major and minor allele information as independent character states. RaxML-NG [23] was used to infer phylogenetic trees. The consensus sequence was analyzed with a GTR+*γ* model, while the PROTGTR+*γ* model was used for the alignment with intra-host diversity and major/minor variant information.

Several metrics were used to compare the 200 inferred phylogenetic trees with their respective starting phylogeny from which the sequences were simulated (Supplementary Table 2). We chose metrics available in R suitable for unrooted trees, using the option ‘rooted=FALSE’ where appropriate. The Robinson-Foulds (RF) distance [37] calculates the number of splits differing between both phylogenetic trees. For the weighted Robinson-Foulds (wRF), the distance is expressed in terms of the branch lengths of the differing splits. The Kuhner-Felsenstein distance [38] considers the edge length differences in all splits, regardless of whether the topology is shared or not. Last, the Penny-Steel distance or path difference metric [39] calculates the pairwise differences in the path of each pair of tips, with the weighted Penny-Steel distance (wPS) using branch length to compute the path differences. All functions were used as implemented in the package *phangorn* [36] within R [35].

### Amplification and whole-genome sequencing

SARS-CoV-2 real-time qPCR confirmed isolates from London hospitals were collected as part of the routine diagnostic service at Great Ormond Street Hospital NHS Foundation Trust (GOSH) [40] and the COVID-19 Genomics UK Consortium (COG-UK) [17] between March and December 2020, in addition to epidemiological and patient metadata (Supplementary Table 1). SARS-CoV-2 whole-genome sequencing was performed by UCL Genomics. cDNA and multiplex PCR reactions were prepared following the ARTIC nCoV-2019 sequencing protocol [41]. The ARTIC V3 primer scheme [42] was used for the multiplex PCR, with a 65°C, 5 min annealing/extension temperature. Pools 1 and 2 multiplex PCRs were run for 35 cycles. 5µL of each PCR were combined and 20µL nuclease-free water added. Libraries were prepared on the Agilent Bravo NGS workstation option B using Illumina DNA prep (Cat. 20018705) with unique dual indexes (Cat. 20027213/14/15/16). Equal volumes of the final libraries were pooled, bead purified and sequenced on the Illumina NextSeq 500 platform using a Mid Output 150 cycle flowcell (Cat. 20024904) (2 x 75bp paired ends) at a final loading concentration of 1.1pM.

### Whole-genome sequence analysis of SARS-CoV-2 sequences

Raw illumina reads were quality trimmed using Trimmomatic [43] with a minimum mean quality per base of 20 in a 4-base wide sliding window. The 5 leading and trailing bases of each read were removed, and reads with an average quality lower than 20 were discarded. The resulting reads were aligned against the Wuhan-Hu-1 reference genome (GenBank NC_45512.2, GISAID EPI_ISL_402125) using BWA-mem v0.7.17 with default parameters [44]. The alignments were subsequently sorted by position using SAMtools v1.14 [45]. Primer sequences were masked using ivar [46].

Single-nucleotide variants were identified using the pileup functionality of samtools [45] via the pysam package in Python (https://github.com/pysam-developers/pysam). Variants were further filtered using bcftools [18]. Only variants with a minimum depth of 50x and a minimum base quality and mapping quality of 30 were kept. Additionally, variants within low complexity regions identified by sdust (https://github.com/lh3/sdust) were removed. For positions where only one base was present, the minimum depth was 20 reads, with at least 5 reads in each direction. Positions with low frequency variants were filtered if the total coverage at that position was less than 100x, with at least 20 reads in total and 5 reads in each strand supporting each of the main two alleles.

Two different alignments were prepared from the data. First, an alignment of the consensus sequence where the most prevalent base at each position was kept. Variants where the most prevalent allele was not supported by more than 60% of the reads were considered ambiguous. Additionally, an alignment reflecting within-sample variation at each position as well as which base is the most prevalent and which one appears at a lower frequency by using the IUPAC nomenclature for amino acids [22].

For the two different alignments, maximum likelihood phylogenies were inferred by using RAxML-NG [23] with 20 starting trees (10 random and 10 parsimony), 100 bootstrap replicates, and a minimum branch length of 10*^−^*^9^. For the consensus sequence, the GTR model was used. For the alignment reflecting within-host diversity, a model with amino acid nomenclature (PROTGTR) was used. All models allowed for a *γ* distributed rate of variation among sites.

## Data availability

Samples sequenced as part of this study have been submitted to the European Nucleotide Archive under accession PRJEB53224. Sample metadata is included in Supplementary Data File 1.

## Code availability

All custom code used in this article can be accessed at https://github.com/arturotorreso/scov2_withinHost.git.

## Acknowledgements

The authors dedicate this article to the hospital staff members and patients who died of coronavirus disease 2019. They also thank all staff and patients who have taken part in the study. In addition, the authors are very grateful to the Great Ormond Street laboratory staff, the staff at the Camelia Botnar Laboratory, the Great Ormond Street Institute of Child Health and the COVID-19 sequencing team at UCLG who worked tirelessly to ensure that all polymerase chain reaction tests and sequencing work were completed in a timely manner during the COVID-19 pandemic. All authors acknowledge UCL Computer Science Technical Support Group (TSG) and the UCL Department of Computer Science High Performance Computing Cluster. LG was supported by the Wellcome Trust (201470/Z/16/Z), the National Institute of Allergy and Infectious Diseases of the National Institutes of Health under award number 1R01AI146338 and by the GOSH/ICH Biomedical Research Centre. XD was supported by the NIHR Health Protection Research Unit in Genomics and Enabling Data.

## Author Contributions

ATO, LG, XD, and MK conceived and designed the study. LG, NS, SR, RW, KH, CW and JB performed and advised on sample preparation and whole-genome sequencing work. JH and HD collected and analyzed epidemiological data. ATO, MK, XD, RG, JB and LG performed and advised on statistical and computational analyses. ATO and LG wrote the manuscript with input from all co-authors. All authors read and approved the final manuscript.

## Ethics declarations

Ethical approval was obtained for all individual studies from which this data was derived.

## Supplementary Information

**Supplementary Table 1:**
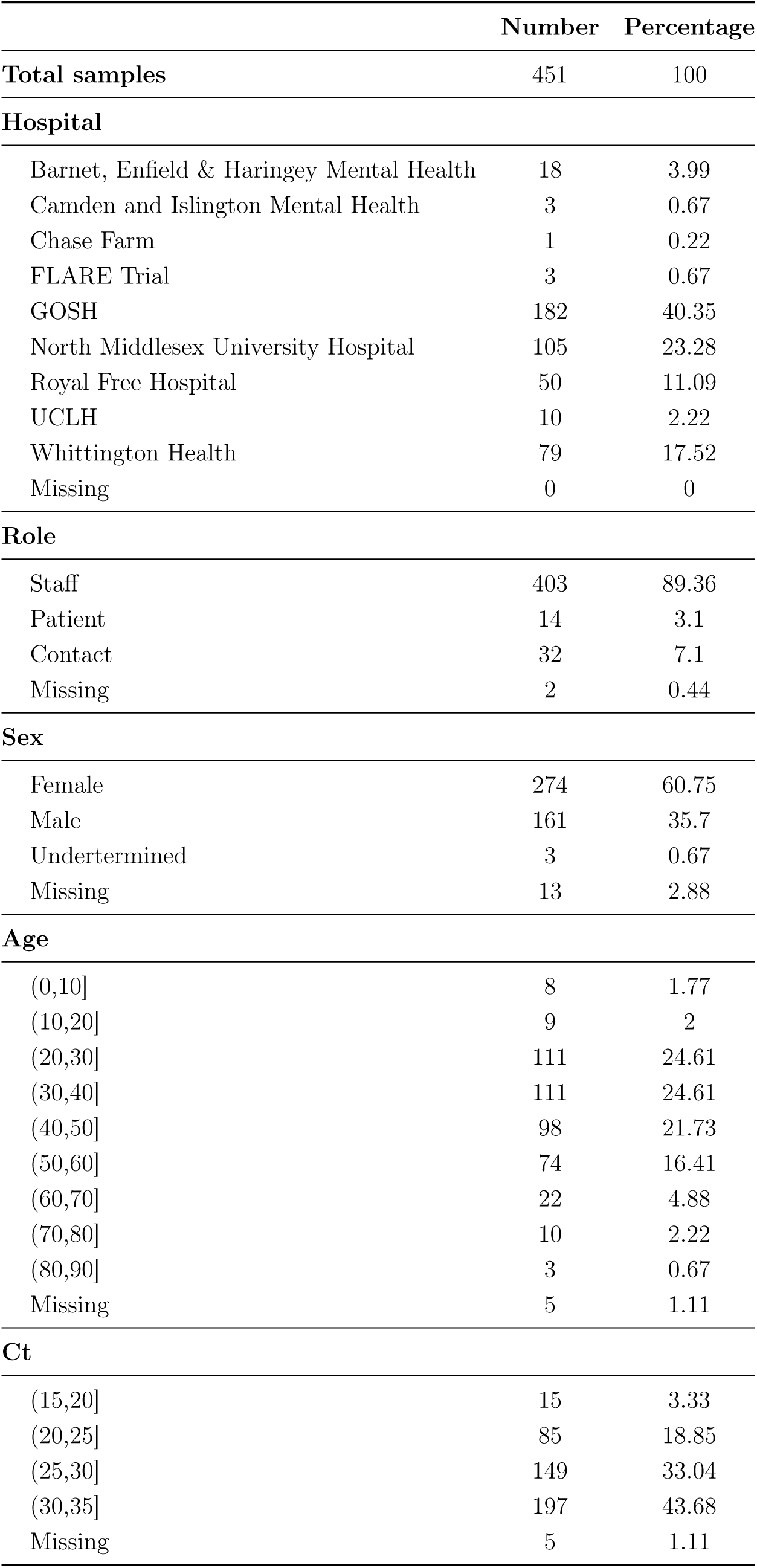
Sample collection and demographics.

**Supplementary Table 2:**
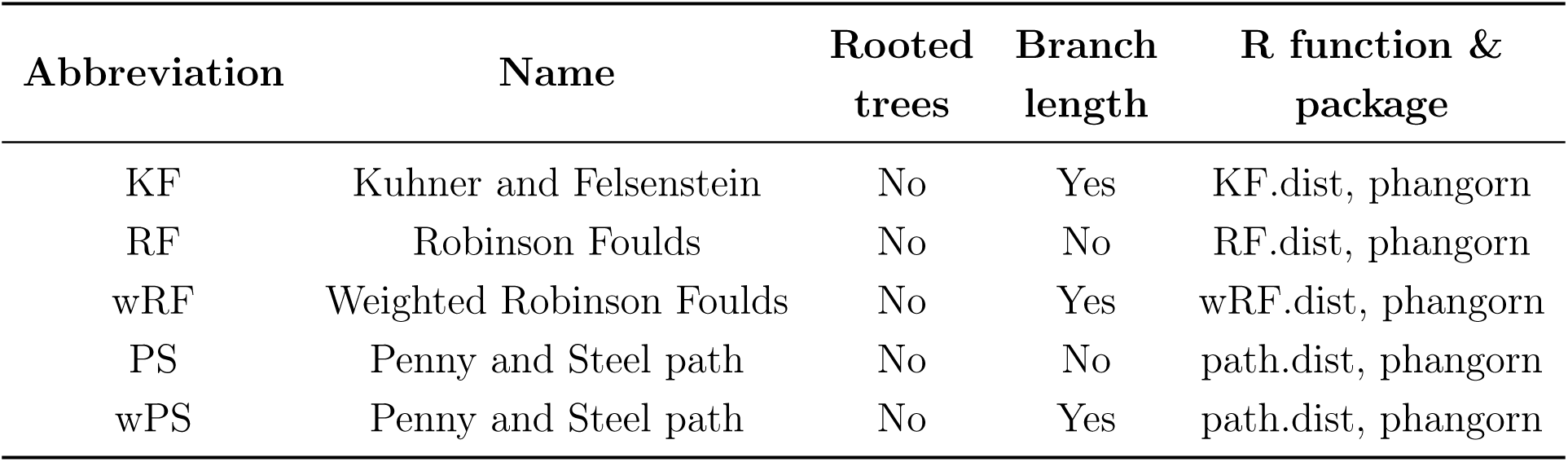
Metrics used for phylogenetic tree comparison.

**Supplementary Table 3:**
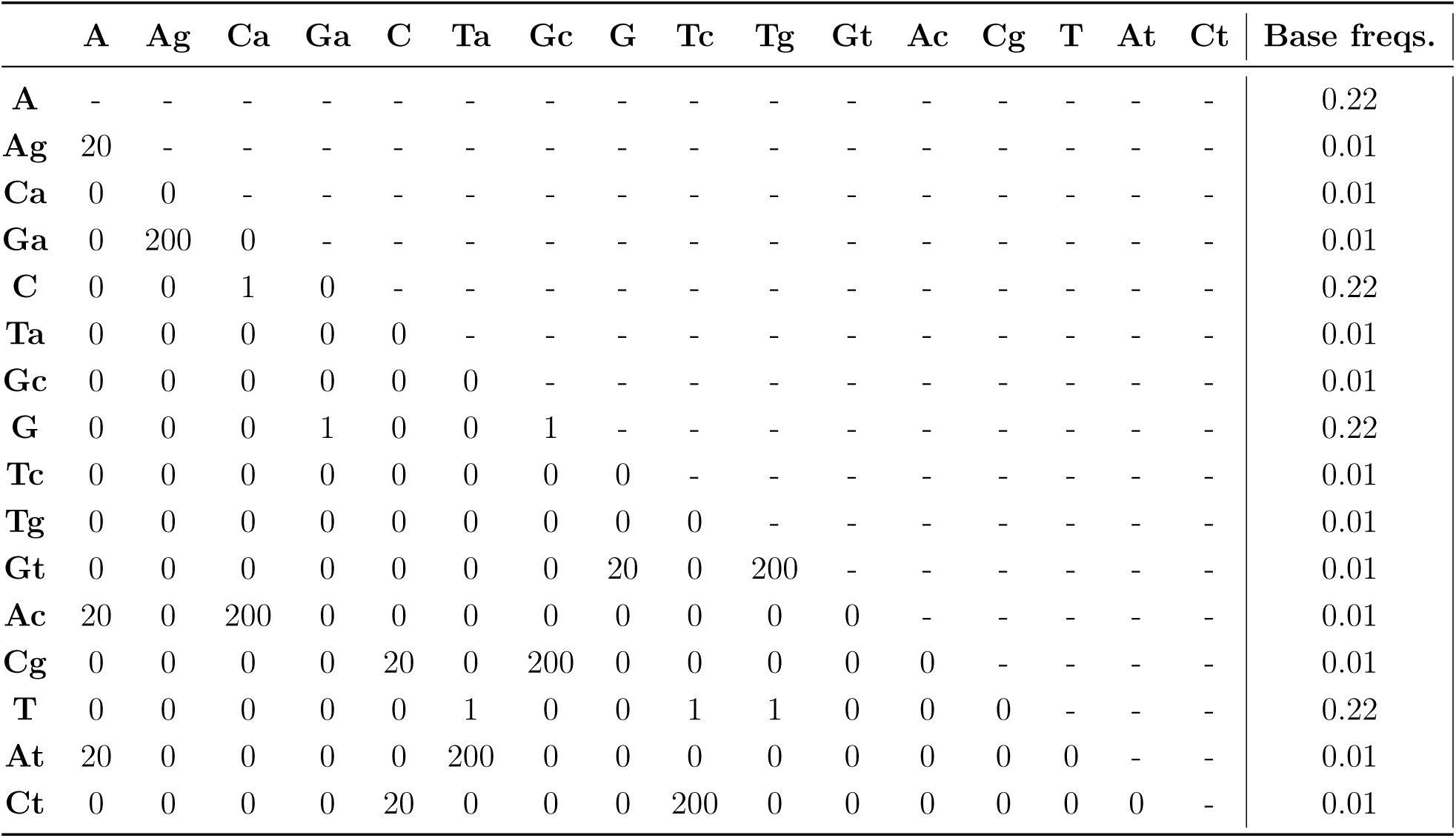
Transition/transversion rates and base frequencies of the known simulated tree.

|  | A | Ag | Ca | Ga | C | Ta | Gc | G | Tc | Tg | Gt | Ac | Cg | T | At | Ct | Base freqs. |
| --- | --- | --- | --- | --- | --- | --- | --- | --- | --- | --- | --- | --- | --- | --- | --- | --- | --- |
| **A** | - | - | - | - | - | - | - | - | - | - | - | - | - | - | - | - | 0.22 |
| **Ag** | 20 | - | - | - | - | - | - | - | - | - | - | - | - | - | - | - | 0.01 |
| **Ca** | 0 | 0 | - | - | - | - | - | - | - | - | - | - | - | - | - | - | 0.01 |
| **Ga** | 0 | 200 | 0 | - | - | - | - | - | - | - | - | - | - | - | - | - | 0.01 |
| **C** | 0 | 0 | 1 | 0 | - | - | - | - | - | - | - | - | - | - | - | - | 0.22 |
| **Ta** | 0 | 0 | 0 | 0 | 0 | - | - | - | - | - | - | - | - | - | - | - | 0.01 |
| **Gc** | 0 | 0 | 0 | 0 | 0 | 0 | - | - | - | - | - | - | - | - | - | - | 0.01 |
| **G** | 0 | 0 | 0 | 1 | 0 | 0 | 1 | - | - | - | - | - | - | - | - | - | 0.22 |
| **Tc** | 0 | 0 | 0 | 0 | 0 | 0 | 0 | 0 | - | - | - | - | - | - | - | - | 0.01 |
| **Tg** | 0 | 0 | 0 | 0 | 0 | 0 | 0 | 0 | 0 | - | - | - | - | - | - | - | 0.01 |
| **Gt** | 0 | 0 | 0 | 0 | 0 | 0 | 0 | 20 | 0 | 200 | - | - | - | - | - | - | 0.01 |
| **Ac** | 20 | 0 | 200 | 0 | 0 | 0 | 0 | 0 | 0 | 0 | 0 | - | - | - | - | - | 0.01 |
| **Cg** | 0 | 0 | 0 | 0 | 20 | 0 | 200 | 0 | 0 | 0 | 0 | 0 | - | - | - | - | 0.01 |
| **T** | 0 | 0 | 0 | 0 | 0 | 1 | 0 | 0 | 1 | 1 | 0 | 0 | 0 | - | - | - | 0.22 |
| **At** | 20 | 0 | 0 | 0 | 0 | 200 | 0 | 0 | 0 | 0 | 0 | 0 | 0 | 0 | - | - | 0.01 |
| **Ct** | 0 | 0 | 0 | 0 | 20 | 0 | 0 | 0 | 200 | 0 | 0 | 0 | 0 | 0 | 0 | - | 0.01 |

**Supplementary Table 4:** Inferred transition/transversion rates and base frequencies when using the consensus sequence. Numbers show the average of 100 simulations.

|  | A | C | G | T | Base freqs |
| --- | --- | --- | --- | --- | --- |
| A | - | - | - | - | 0.255 |
| C | 0.974 | - | - | - | 0.250 |
| G | 1.088 | 1.148 | - | - | 0.247 |
| T | 0.854 | 0.996 | 1 | - | 0.248 |

**Supplementary Table 5:** Inferred transition/transversion rates and base frequencies when accounting for within-host diversity. Numbers show the average of 100 simulations

|  | A | Ag | Ca | Ga | C | Ta | Gc | G | Tc | Tg | Gt | Ac | Cg | T | At | Ct | Base freqs. |
| --- | --- | --- | --- | --- | --- | --- | --- | --- | --- | --- | --- | --- | --- | --- | --- | --- | --- |
| A | - | - | - | - | - | - | - | - | - | - | - | - | - | - | - | - | 0.22 |
| Ag | 17.35 | - | - | - | - | - | - | - | - | - | - | - | - | - | - | - | 0.01 |
| Ca | 0 | 0 | - | - | - | - | - | - | - | - | - | - | - | - | - | - | 0.01 |
| Ga | 0 | 185.96 | 0 | - | - | - | - | - | - | - | - | - | - | - | - | - | 0.01 |
| C | 0 | 0 | 1.86 | 0 | - | - | - | - | - | - | - | - | - | - | - | - | 0.22 |
| Ta | 0 | 0 | 0 | 0 | 0 | - | - | - | - | - | - | - | - | - | - | - | 0.01 |
| Gc | 0 | 0 | 0 | 0 | 0 | 0 | - | - | - | - | - | - | - | - | - | - | 0.01 |
| G | 0 | 0 | 0 | 1.16 | 0 | 0 | 1.08 | - | - | - | - | - | - | - | - | - | 0.21 |
| Tc | 0 | 0 | 0 | 0 | 0 | 0 | 0 | 0 | - | - | - | - | - | - | - | - | 0.01 |
| Tg | 0 | 0 | 0 | 0 | 0 | 0 | 0 | 0 | 0 | - | - | - | - | - | - | - | 0.01 |
| Gt | 0 | 0 | 0 | 0 | 0 | 0 | 0 | 21.64 | 0 | 196.56 | - | - | - | - | - | - | 0.01 |
| Ac | 16.51 | 0 | 183.53 | 0 | 0 | 0 | 0 | 0 | 0 | 0 | 0 | - | - | - | - | - | 0.01 |
| Cg | 0 | 0 | 0 | 0 | 21.20 | 0 | 200 | 0 | 0 | 0 | 0 | 0 | - | - | - | - | 0.01 |
| T | 0 | 0 | 0 | 0 | 0 | 0.98 | 0 | 0 | 1.05 | 1.26 | 0 | 0 | 0 | - | - | - | 0.21 |
| At | 17.70 | 0 | 0 | 0 | 0 | 175.58 | 0 | 0 | 0 | 0 | 0 | 0 | 0 | 0 | - | - | 0.01 |
| Ct | 0 | 0 | 0 | 0 | 20.09 | 0 | 0 | 0 | 188.86 | 0 | 0 | 0 | 0 | 0 | 0 | - | 0.01 |

**Supplementary Figure 1:**
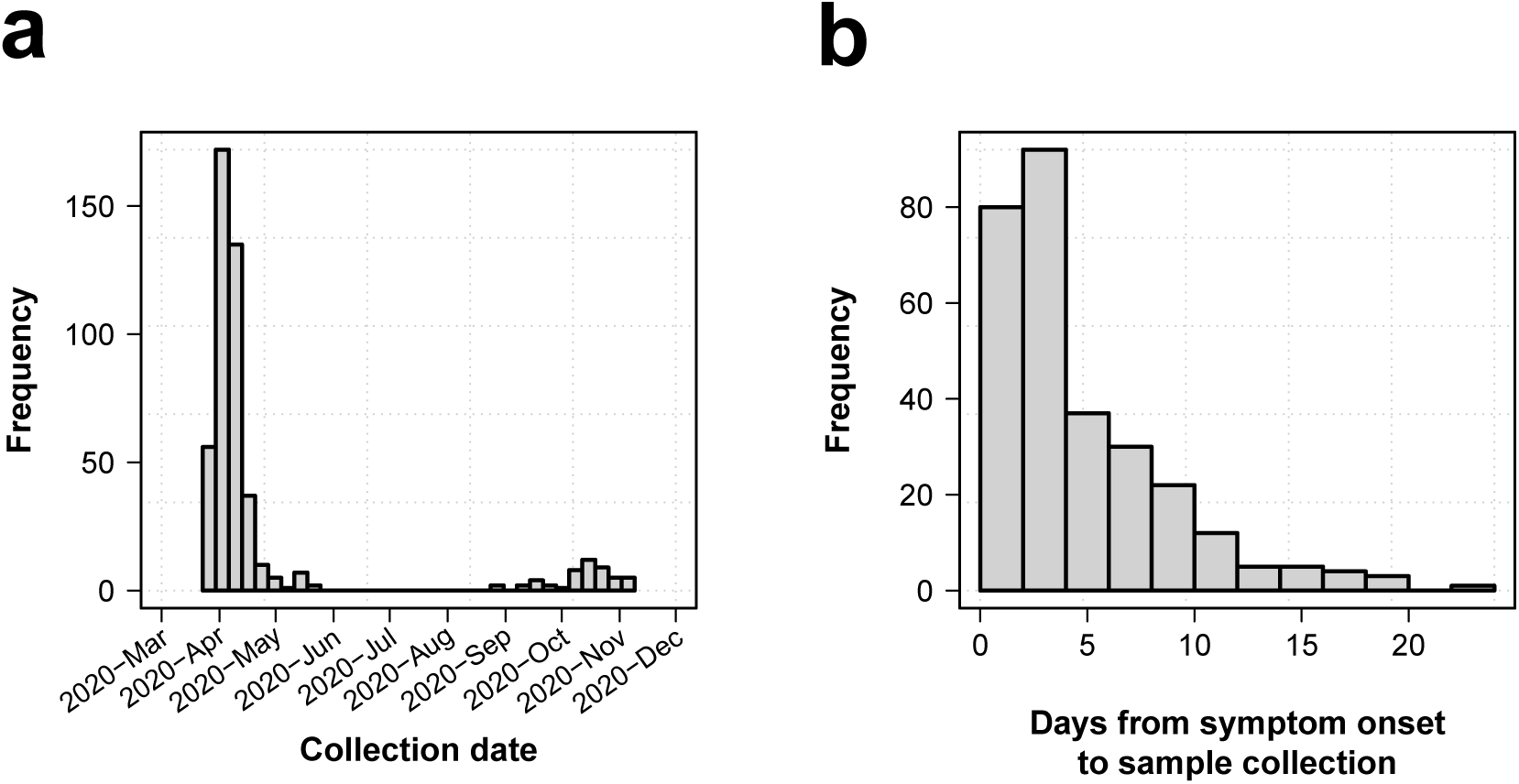
Collection date distribution and time from symptom and days from symptom onset. (a) Distribution of collection dates. (b) Histogram of time from symptom onset to sample collection.

**Supplementary Figure 2:**
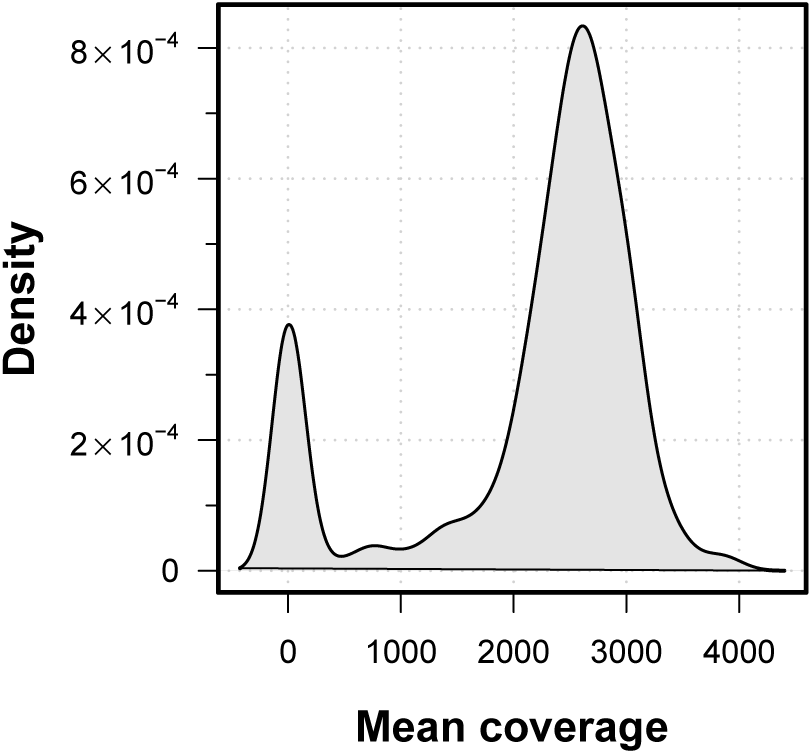
Sample mean coverage distribution. Density distribution of mean coverage.

**Supplementary Figure 3:**
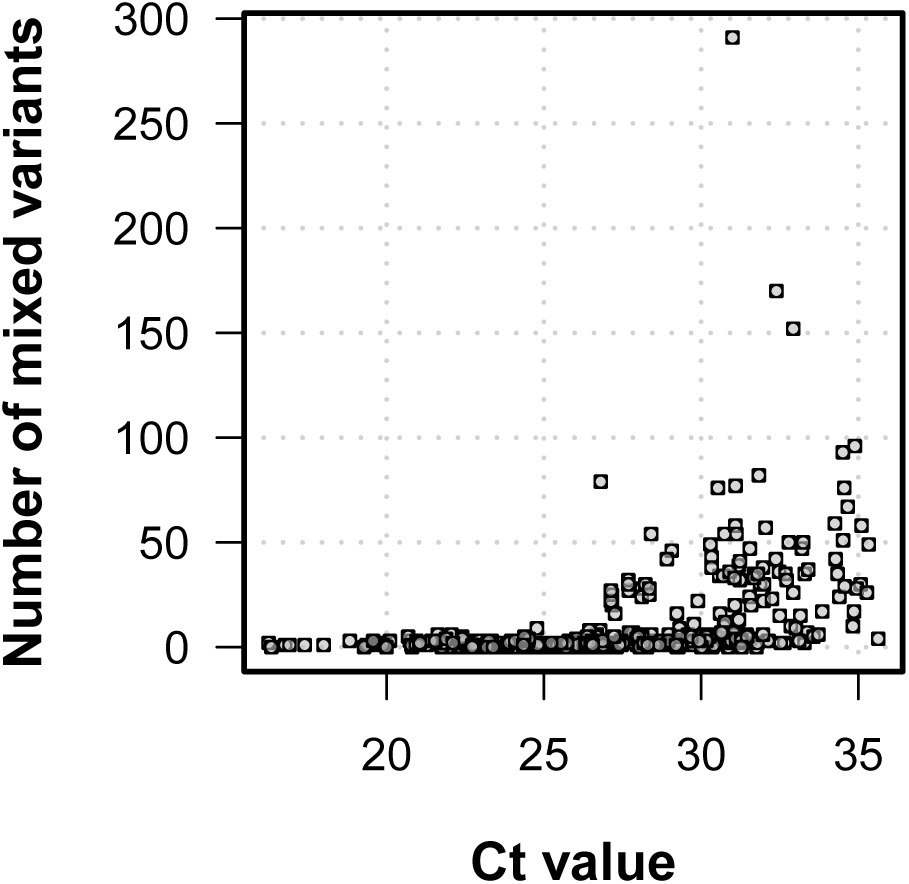
Number of low frequency variants and C*_t_* value. Higher C*_t_* values were linked to a higher number of within-sample variation.

**Supplementary Figure 4:**
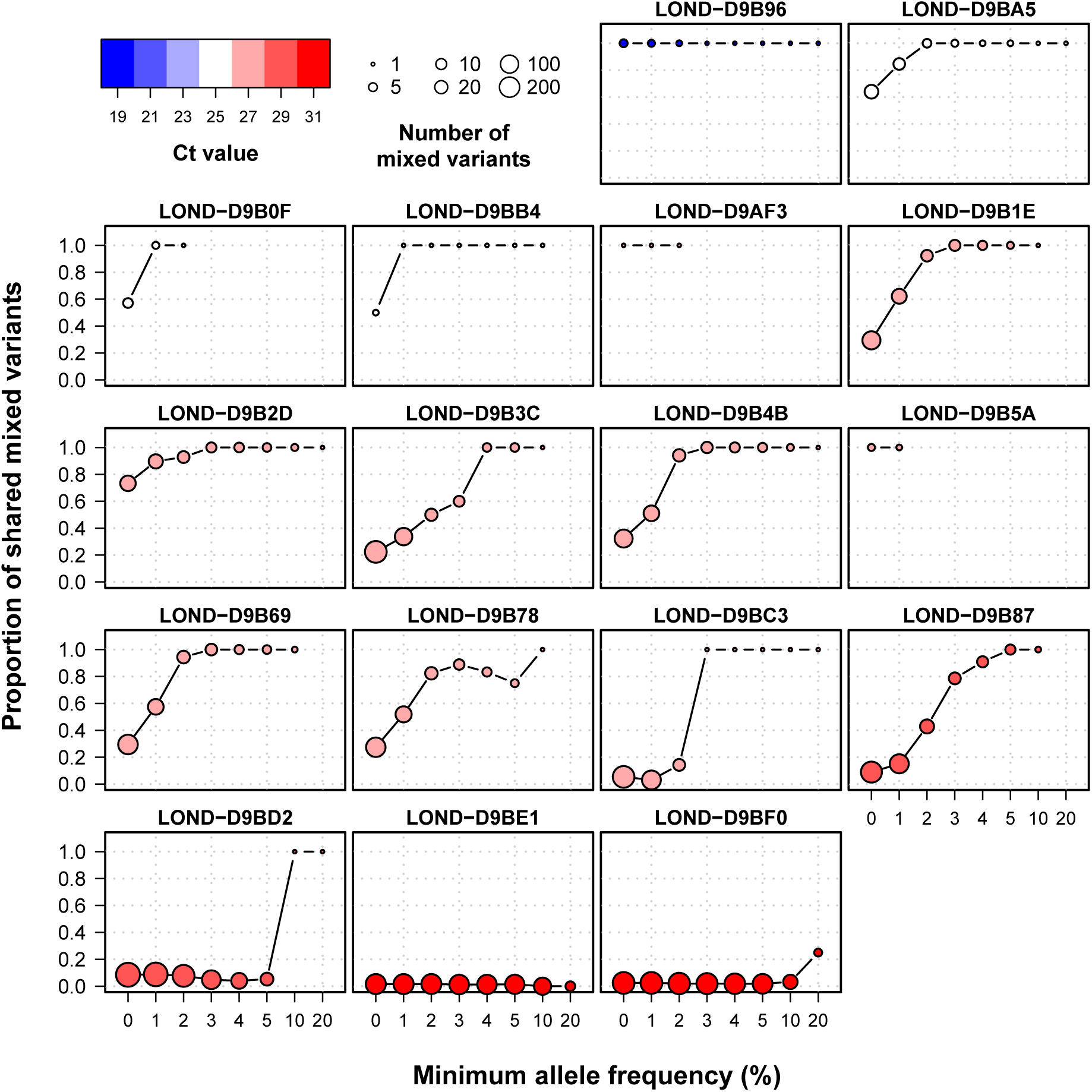
Proportion of shared mixed variants between duplicated samples using different filters of allele frequency.Individual plots of shared within-host variants between technical duplicates using increasing thresholds of allele frequency. Colors represent C*_t_* value, while the size of the point shows the total number of within-host variants between the two samples.

**Supplementary Figure 5:**
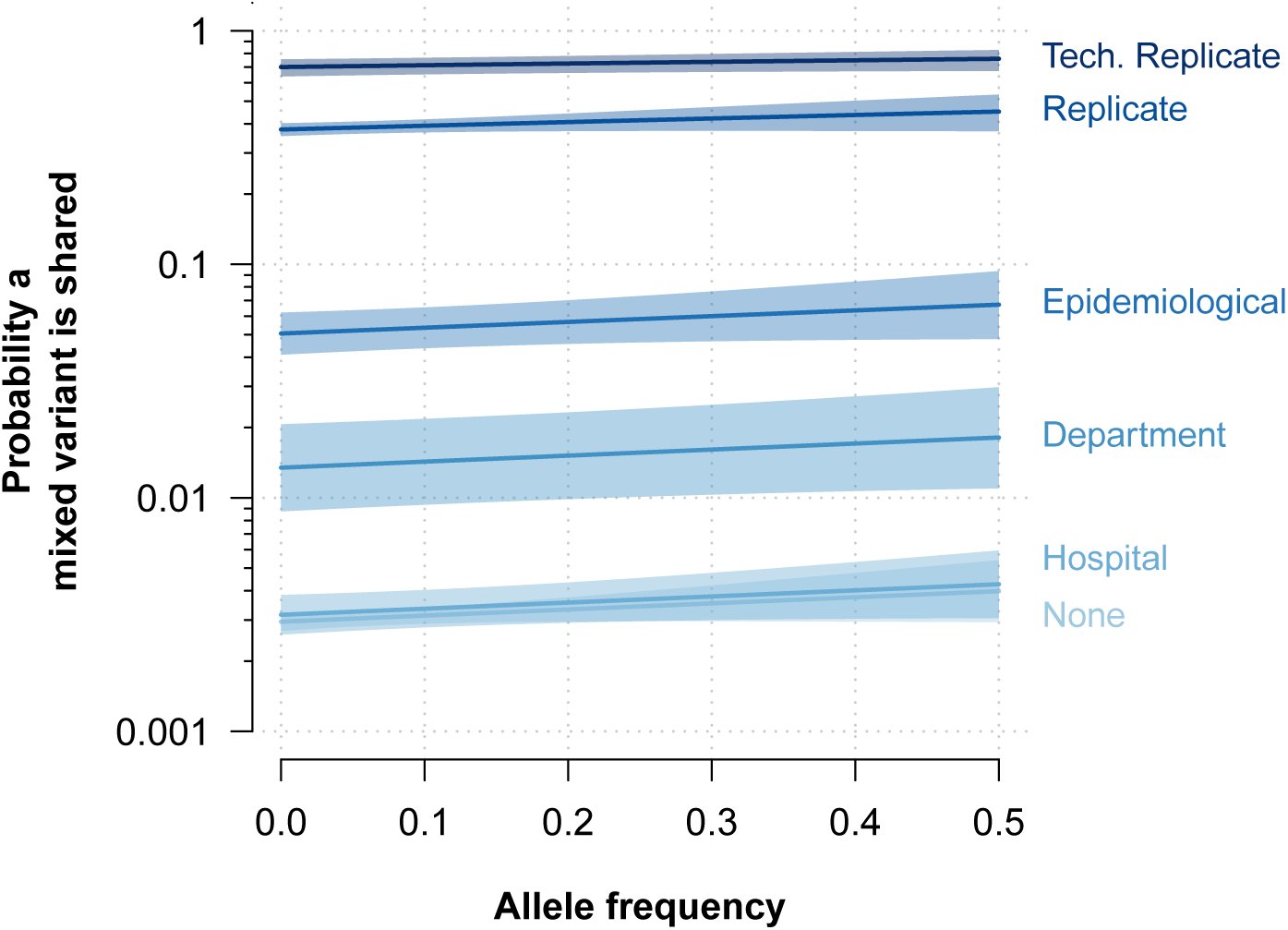
Probability that mixed variants are shared. Probability that low frequency variants are shared inferred with a logistic model with allele frequency and epidemiological relationship as independent variable and whether a variant is shared or not as dependent variable. Y-axis in logarithmic scale for representation.

**Supplementary Figure 6:**
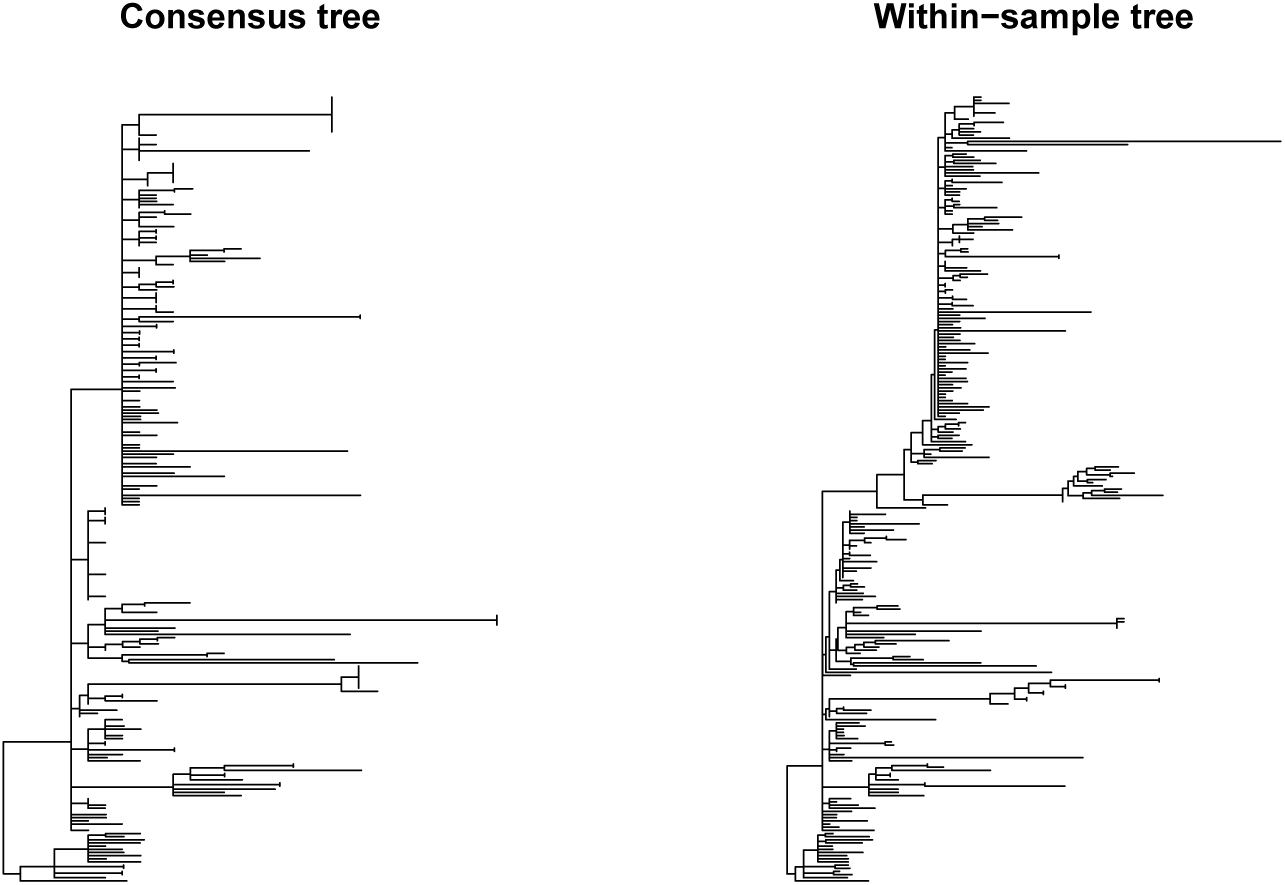
Phylogenetic trees for SARS-CoV-2. SARS-CoV-2 phylogenetic trees inferred from consensus sequences (left) and an alignment with major and minor variant information (right) .

**Supplementary Figure 7:**
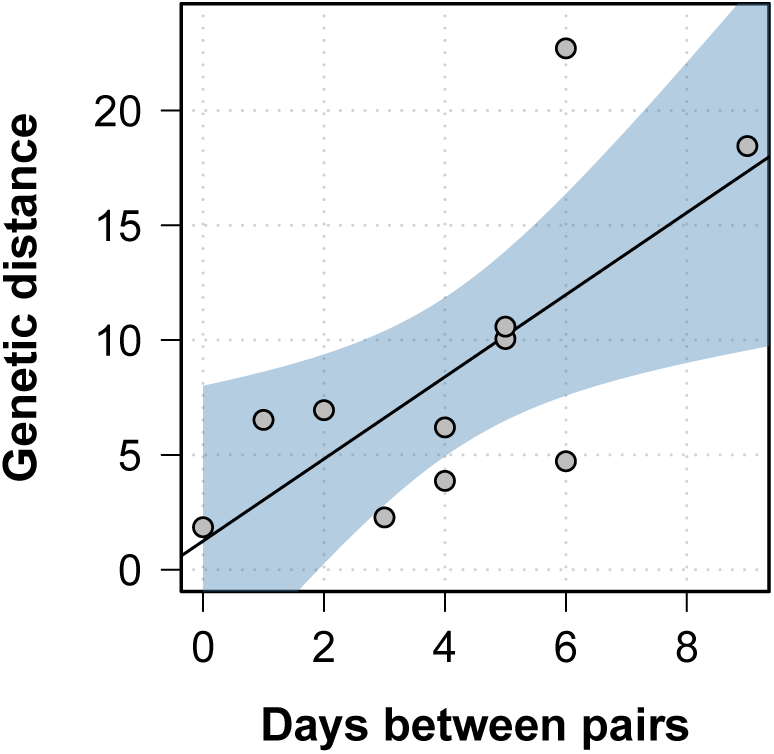
Genetic distance between longitudinal samples. The genetic distance in the phylogenetic tree inferred using within-sample diversity increased as the between longitudinal samples progressed. Black line shows the best fit in a linear model, while the blue shaded area represents the 95% CI.

**Supplementary Figure 8:**
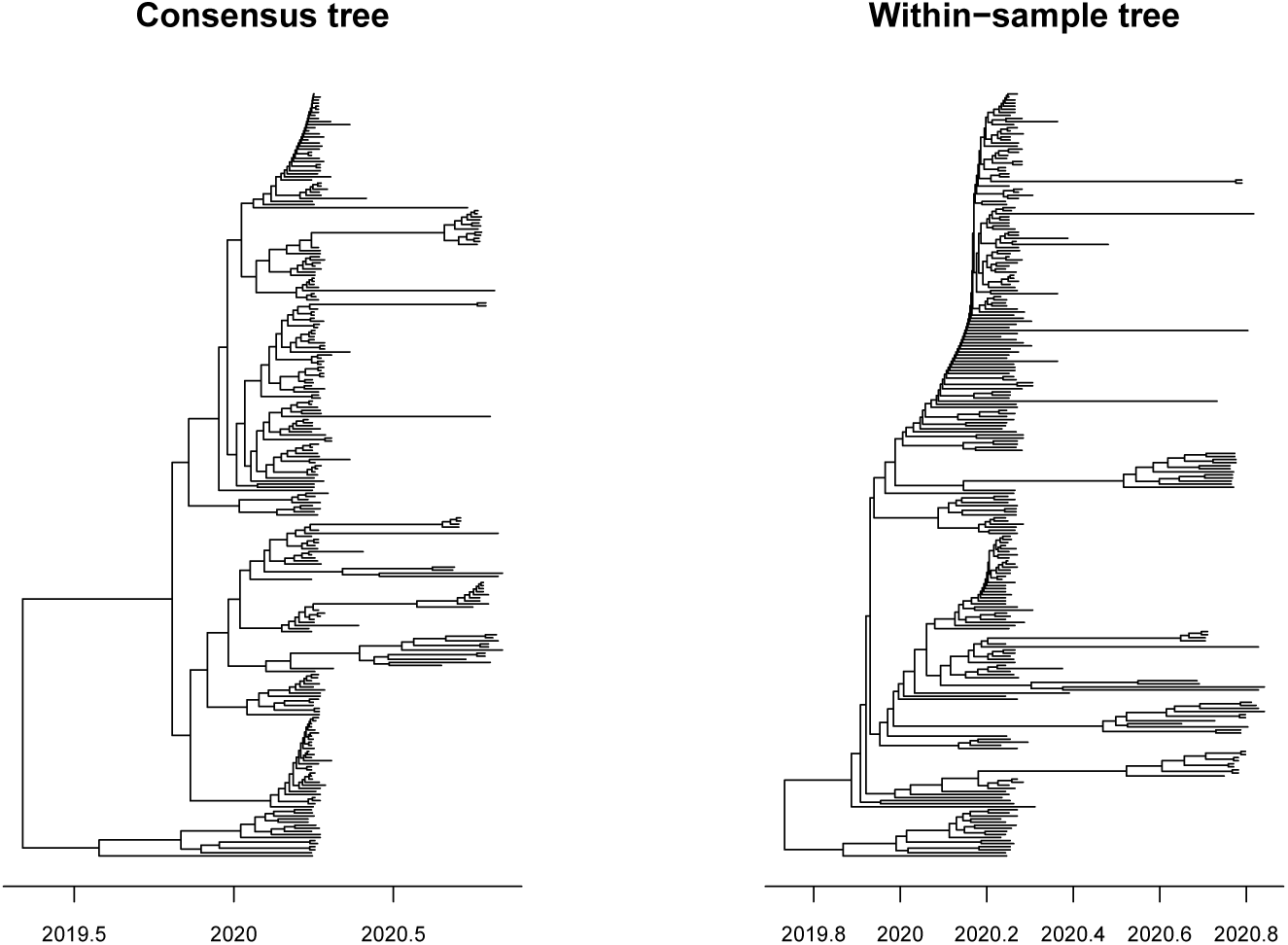
Time calibrated phylogenetic trees for SARS-CoV-2. SARS-CoV-2 phylogenetic trees inferred from consensus sequences (left) and an alignment with major and minor variant information (right). Branch lengths are measured in years.

**Supplementary Figure 9:**
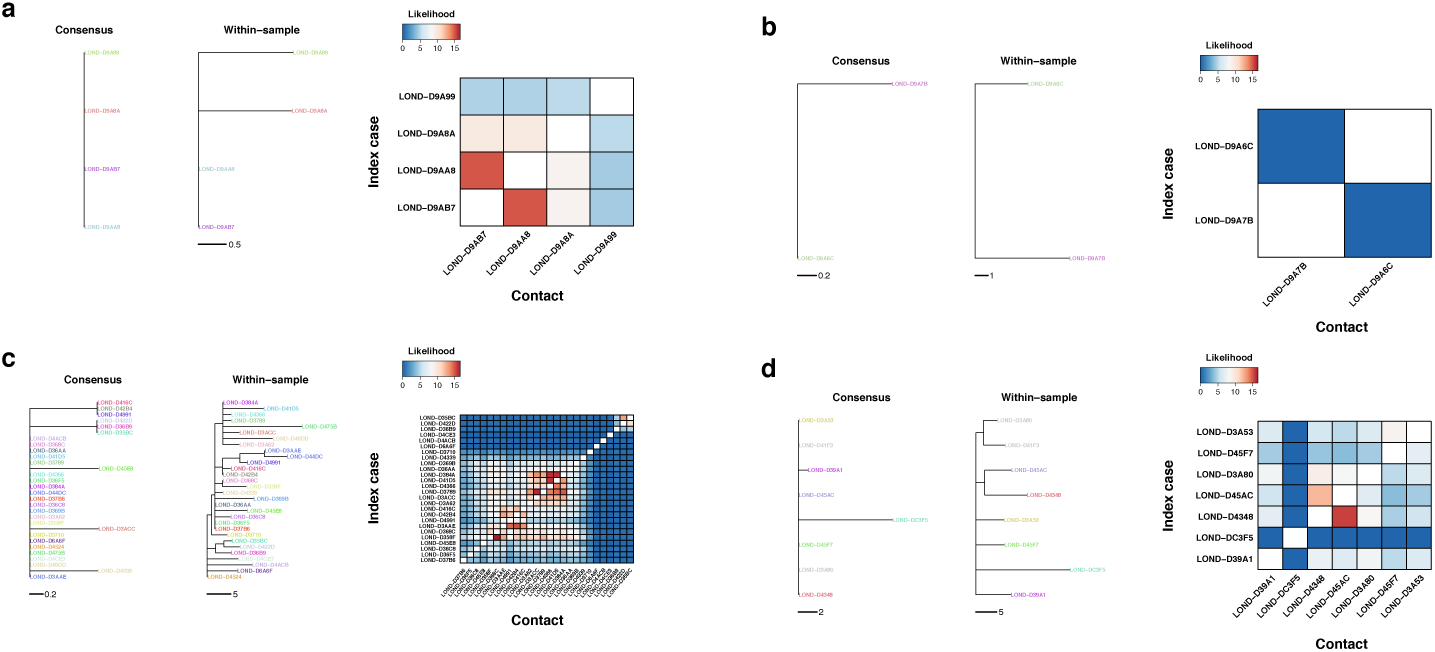
Phylogenetic and transmission for SARS-CoV- 2 outbreaks. a-d Phylogenies of SARS-CoV-2 outbreaks. The branch lengths are in units of substitutions per genome, and the scales are shown under the trees. Colors represent samples from the same individual. Samples with the same name are technical replicates. Left tree of each panel shows the phylogeny inferred with the consensus alignment. Right tree represents the phylogeny inferred using within-sample variation. Heatmap shows the likelihood of direct transmission for each pair of samples in a SEIR model of transmission. Vertical axis is the infector while the horizontal axis shows the infectee.

